# PIONEER: A Periplasmic Display Platform for Synthetic Biology-Based Screening of Genetically Encoded Protein Regulators

**DOI:** 10.1101/2025.10.26.684622

**Authors:** Jacob B. Rowe, Kyutae Lee, Daniel G. Isom

**Affiliations:** Department of Molecular and Cellular Pharmacology, University of Miami Miller School of Medicine, Miami, FL, USA; Tumor Biology Program, University of Miami Sylvester Comprehensive Cancer Center, Miami, FL, USA; Frost Institute for Data Science and Computing, Miami, FL, USA; Frost Institute for Chemistry and Molecular Sciences, Miami, FL, USA

**Keywords:** GPCRs, DCyFIR, SynBio, genetically encoded modulators, nanobodies, yeast

## Abstract

Periplasmic display in yeast is a promising but underdeveloped method for screening and studying genetically encoded biomolecules. We present PIONEER, a modular platform that localizes peptides, proteins, and nanobodies to the membrane-proximal periplasmic space of *S. cerevisiae*, enabling direct interrogation of membrane protein function. By optimizing secretion signals and display scaffolds, PIONEER ensures stable ligand retention and robust autocrine signaling. The ability to assess peptide and protein ligand activity through signaling, rather than binding alone, marks a key advance toward function-first screening. Applied to human G protein-coupled receptors (GPCRs), the system enables detection of surface expression, ligand activity profiling, and classification of nanobody regulators, including antagonists and conformational stabilizers. It also distinguishes intrabodies based on their functional effects and chaperone activity. Although demonstrated with GPCRs, PIONEER is adaptable to a wide range of membrane and soluble protein targets. As AI-driven design expands the space of candidate ligands and binders, this platform offers a scalable method for linking predicted sequences and their structures to biological function.

## Introduction

Yeast is a versatile eukaryotic platform widely used in synthetic biology for applications ranging from the development and production of biopharmaceuticals^1^, biofuels^2^, sustainable materials^3^, and natural products^4–6^. For example, yeast has been engineered to synthesize human insulin^7^, the anti-malarial drug artemisinin^8^, and complex proteins requiring post-translational modifications like glycosylation and disulfide bond formation^1^. Yeast systems are also used to design genetic circuits, enabling precise control of gene expression^9^, and as biosensors for environmental or medical diagnostics^10^. Furthermore, their industrial scalability makes them invaluable for large-scale production processes^11^.

Yeast display is a foundational synthetic biotechnology that uses engineered yeast, typically *S. cerevisiae*, to present proteins and peptides on the cell surface. The protein of interest is genetically fused to Aga2p, which is secreted and forms disulfide bonds with the cell wall-anchored Aga1p, tethering the target protein to the cell exterior^12^. This configuration makes displayed proteins accessible for binding assays, high-throughput screening, and interaction analysis. As such, this technique is widely used for antibody affinity maturation through directed evolution^13,14^, nanobody discovery^15,16^, enzyme engineering^17^, studying protein-protein interactions^18^, and biosensor design^19^.

In contrast, yeast periplasmic display is conceptually the inverse of surface display and remains vastly underexplored. It offers unique advantages, particularly for applications involving transmembrane proteins. This method localizes recombinant proteins to the space between the plasma membrane and the inner cell wall, a confined periplasmic-like compartment. Proteins fused to anchors such as invertase (Suc2p) or engineered Flo1p variants are secreted from the cytoplasm but retained near the membrane^20–22^, enabling direct interaction with membrane-embedded targets.

Unlike surface display, which primarily supports the identification of binding partners, periplasmic display enables functional selection (Fig. S1). This allows for the discovery of genetically encoded peptides and proteins that not only bind but also modulate transmembrane protein activity through agonism, inhibition, or allosteric regulation. Because each ligand is barcoded in the yeast genome or on a plasmid, there is a one-to-one correspondence between function and sequence, simplifying downstream screening applications and identification.

Here, we present PIONEER (**P**eriplasmic d**I**splay **O**f ge**N**etically **E**ncoded r**E**gulato**R**s), an advanced platform for discovering and characterizing genetically encoded modulators of G protein-coupled receptor (GPCR) signaling. Built on our previously established DCyFIR platform^21,23–25^, PIONEER employs a humanized yeast mating pathway along with optimized secretion signals and cell wall display proteins to support studies of peptide- and protein-based ligands. We demonstrate its utility for detecting GPCR surface expression, profiling ligand-activated signaling, and characterizing GPCR-targeting nanobodies. PIONEER provides a powerful tool to accelerate the development of next-generation GPCR modulators for structure-based drug discovery, basic research, therapeutic development, and theragnostic applications.

## Results

To optimize periplasmic display, we developed a modular platform in *S. cerevisiae*, using humanized GPCR signaling as a proof of concept. As shown in Fig. 1a, the platform comprises three key components. First, an efficient secretion signal directs proteins to the periplasmic space. Second, a genetically encoded protein ligand serves as the functional input. Third, a cell wall display domain retains the ligand in the periplasm, presents it on the inner cell wall, and preserves access to membrane-localized receptors.

**Fig. 1.**
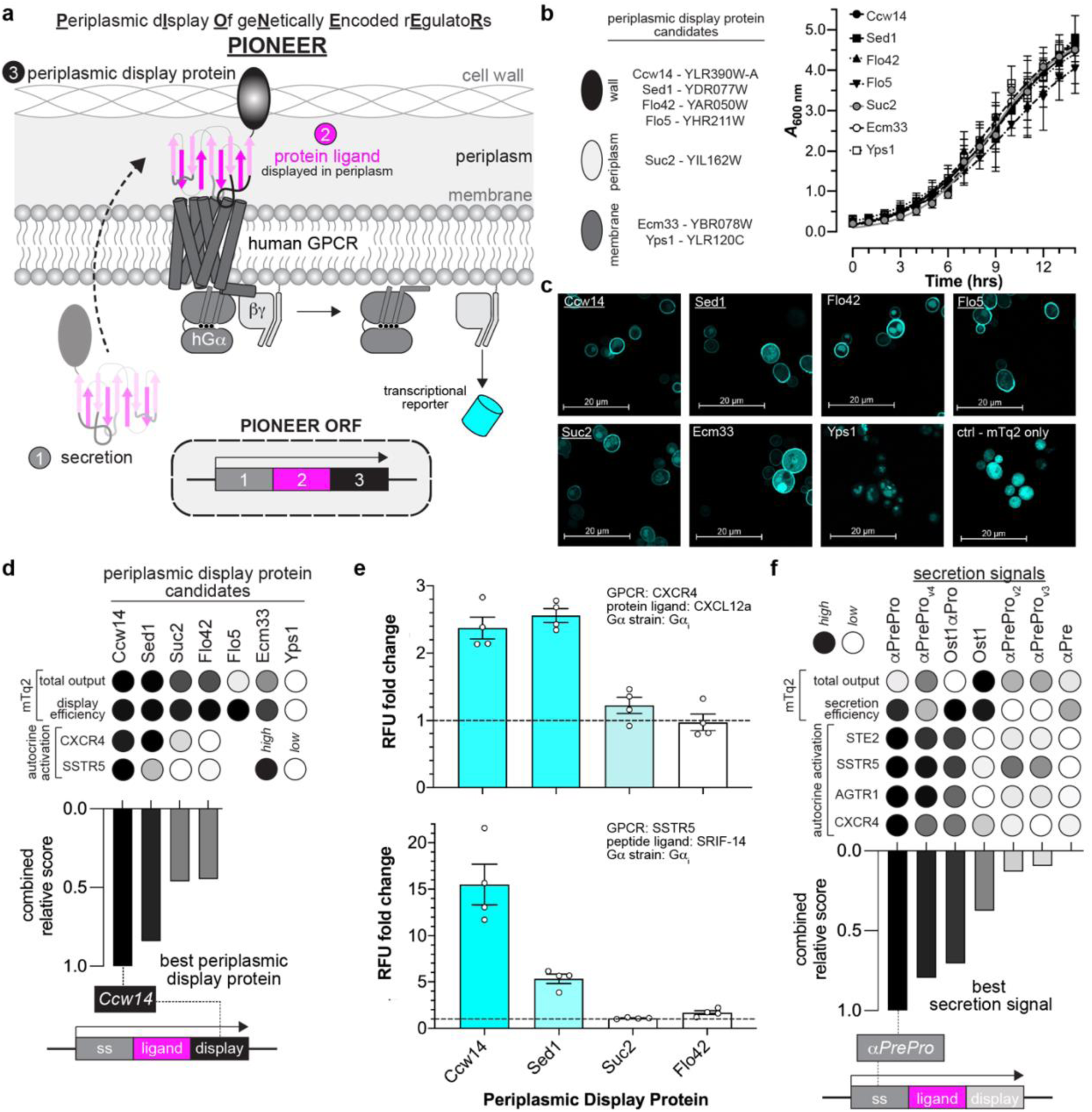
Design and validation of the PIONEER platform. **a**, Schematic of the PIONEER system. Human GPCR signaling in the DCyFIR system is modulated by periplasmically displayed proteins and detected via transcriptional activation of a fluorescent reporter. **b**, Effects of display protein overexpression on yeast growth, assessed by optical density at 600 nm (A₆₀₀). All constructs included the αPre secretion signal for comparison. **c**, Confocal microscopy of mTq2-tagged display proteins. Underlining indicates proteins with prominent peripheral localization. **d**, Performance scoring of display proteins in mTq2-tagging and ligand-display assays. **e**, GPCR activation in response to periplasmically displayed ligands in the DCyFIR system, measured in relative fluorescence units (RFU). Fold change is calculated relative to signaling elicited by secreted-only ligands. **f**, Performance scoring of secretion signals in mTq2-tagging and ligand-secretion assays. Data in **b-f** represent mean ± s.d. of *n* = 4 or 8 biological replicates (as indicated for each panel). Ligand-based assays (**d-f**) were performed in a DCyFIR G_αi_ strain, except for *STE2*, which was tested in the parental yeast strain expressing native G_α_ (*Gpa1*). Rankings were normalized to the best-and worst-performing candidates in each assay.

While our demonstration centers on GPCR signaling and builds on the DCyFIR platform, we aimed to design a secretion signal and display domain applicable across synthetic biology. Flo42, the current standard for periplasmic display, is a truncated, GPI-anchored form of the yeast Flo1 protein but yields weak receptor activation compared to exogenous ligand addition^21^. This limited performance may stem from low ligand-Flo42 expression or secretion, or structural interference in the fusion constructs. To improve display system performance, we focused on identifying and validating next-generation display domains.

### Ccw14, Sed1, Suc2, and Flo5 are effective periplasmic display scaffolds

We began by surveying the literature and selecting seven periplasmic display proteins for testing based on relevance and feasibility (Fig. S2a). This set comprised proteins that reversibly bind the cell wall (Ccw14, Sed1, Flo5), diffuse within the periplasm (Suc2), or anchor to the cell membrane (Ecm33, Yps1). We included Flo42 as a benchmark. As shown in Fig. 1b, all yeast strains overexpressing a candidate reached stationary phase within 12 hours, with final optical densities at 600 nm ranging from 3.5 to 4.5. These results indicated that overexpression did not impair growth or cause detectable toxicity.

Next, we examined the localization of each display candidate using mTurquoise2 (mTq2), a bright cyan fluorescent protein, as a test cargo. To ensure consistent targeting to the secretory pathway, we replaced each candidate’s native secretion signal with αPre, a 19-amino acid signal derived from the yeast α-factor^26^, a secreted peptide hormone that triggers mating responses. This yielded a set of αPre-mTq2-*X* fusion constructs, where *X* was the display candidate. We cloned each construct into expression plasmids, introduced them into yeast, and visualized localization using confocal microscopy.

As shown in Fig. 1c, Ccw14, Sed1, Suc2, and Flo5 formed distinct mTq2 fluorescent rings around each yeast cell, with minimal signal in the cytoplasm and organelles. This pattern indicated efficient entry into and transit through the endoplasmic reticulum and Golgi. In contrast, Flo42, Ecm33, and Yps1 showed significant mTq2 fluorescence in vacuoles, suggesting misrouting and degradation. This likely explains the weak signaling observed in our prior studies using Flo42-trapped GPCR agonists^21^. As expected, non-secreted mTq2 accumulated in the cytoplasm and produced bright, diffuse fluorescence. Notably, the fluorescence intensity in Fig. 1c does not reflect total mTq2 levels, as laser power was adjusted to optimize image clarity. Based on localization patterns, we advanced Ccw14, Sed1, Suc2, and Flo5 for further testing.

### Periplasmic display domains effectively prevent detectable cargo escape

Next, we quantified the cell wall display efficiency of our top candidates using disqualified constructs as controls. First, we measured total mTq2 levels in yeast cultures (Fig. S2b). After pelleting, washing, and resuspending the cells, we measured retained mTq2 and calculated display efficiency as the percentage retained (Eq. 1; Fig. S2c). As shown in Fig. 1d, Yps1’s trapping efficiency was set to zero since it was not displayed (Fig. 1c). Flo5, despite its promising localization pattern, showed mTq2 output as low as Yps1 and was disqualified (Fig. S2b). All other candidates, including the controls, exhibited high display efficiency, indicating full mTq2 retention. These results demonstrate that our display domains effectively prevent detectable cargo escape.

### Ccw14 is the most effective scaffold for periplasmic ligand display

Compared to yeast surface display, periplasmic display remains underexplored in synthetic biology, with few studies exploiting protein retention between the plasma membrane and cell wall. Successful signaling applications require constructs that fold into active conformations, minimize steric hindrance of ligand-target interactions, and remain accessible to the extracellular face of transmembrane proteins. To define the scope and power of this approach, we used DCyFIR to screen synthetic autocrine agonism of human somatostatin receptor 5 (SSTR5) and chemokine receptor 4 (CXCR4) with Ccw14, Sed1, and Suc2 ligand-display fusions. Our results establish periplasmic display as a versatile platform for synthetic signaling, while revealing key parameters that govern ligand compatibility and scaffold selection.

As shown in Figs. 1d-e and S3, genetically encoded αPre-display fusions enabled autocrine activation of CXCR4 and SSTR5, which recognize the protein ligand CXCL12a and peptide ligand SRIF-14, respectively. Ccw14 fusions performed best, driving strong activation for both receptors and yielding the highest combined relative score, reflecting total mTq2 output and signaling strength. Sed1 matched Ccw14 for CXCR4 but reached only ~30% of its activity for SSTR5. Suc2 performed poorly, with activation comparable to the Flo42 control and less than half of Ccw14. These results established Ccw14 as the most effective scaffold for periplasmic ligand display.

For CXCR4 and SSTR5, displayed ligands engage the orthosteric site through N-terminal interactions, enabling compatibility with C-terminal Ccw14 fusions. In contrast, ligands that bind via their C-terminus may not be compatible with this design. Testing additional receptors showed that angiotensin receptor 1 (AGTR1) and the C5a receptor (C5aR) fail to activate when fused to Ccw14. Structural studies explain this incompatibility by showing that AGTR1, a key regulator of blood pressure and target of antihypertensive drugs, binds angiotensin II (Ang-II) in a C-terminal-first orientation, with the C-terminal phenylalanine (Phe8) inserting deep into the orthosteric pocket to drive activation^27^. Similarly, C5a engages C5aR by inserting its C-terminal tail into the transmembrane pocket, triggering conformational changes that couple G proteins^28^. In both cases, a C-terminal Ccw14 fusion would occlude critical interactions and prevent activation.

### αPrePro is the optimal secretion signal peptide for Ccw14 display

Having identified Ccw14 as the best display protein, we next optimized the signal peptide responsible for ligand secretion and signaling. Two secretion signals from *S. cerevisiae* are commonly used in yeast biotechnology: αPrePro, which precedes the 13-amino-acid mating pheromone α-factor^26^, and the 22-amino-acid N-terminal signal from Ost1, the α subunit of the oligosaccharyltransferase complex^29^ (Fig. S4a). αPrePro is a bipartite signal comprising a 19-amino-acid αPre segment that directs posttranslational translocation into the ER and a 66-amino-acid Pro region that enables enzymatic processing and Golgi trafficking^30^. In contrast, the Ost1 signal promotes cotranslational translocation into the rough ER and typically yields more efficient secretory entry^31^. To improve displayed ligand secretion, we tested several αPrePro and Ost1 combinations.

As shown in Figs. 1f and S4b-c, we evaluated seven *X*-mTq2 fusion constructs, where *X* denotes the secretion signal. These included wild-type αPrePro and Ost1, the truncated αPre segment, three αPrePro variants (V2, V3, V4) with reported enhancements in Pro-region processing, and a hybrid Ost1αPro sequence previously shown to boost secretion in *P. pastoris*^30^. For each construct, we measured total mTq2 output and secretion efficiency. Ost1 produced the highest output, followed by αPreProV4, V3, and V2, with αPre and αPrePro performing similarly. Ost1αPro yielded the lowest output. In contrast, secretion efficiency was highest for αPrePro, Ost1, and Ost1αPro, followed by αPre and αPreProV4, with V2 and V3 being the least efficient. Based on these results, we selected αPrePro, Ost1, and Ost1αPro as lead candidates.

We next evaluated which secretion signals best supported autocrine GPCR activation in the DCyFIR system. Each secreted agonist functions as a pro-ligand that must undergo enzymatic maturation to remove its N-terminal secretion signal and reveal the active form. To test this, we constructed a panel of *X*-agonist fusions, where *X* denotes the secretion signal being tested. As shown in Figs. 1f and S4d-f, these included *X*-α-factor for Ste2, *X*-SRIF-14 for SSTR5, *X*-CXCL12a for CXCR4, and *X*-AGT-II for AGTR1. Among the secretion signals tested, the native αPrePro consistently supported the strongest GPCR activation, followed by αPreProV4 and Ost1αPro. These results suggest that αPrePro undergoes more robust maturation at steady state, potentially due to parallel processing by dipeptidyl aminopeptidases such as Ste13^26^ and Kex2^32^, as well as the signal peptidase complex^33^, which target conserved -X-Ala-motifs.

Considering total mTq2 output, secretion efficiency, and autocrine activation, we selected αPrePro as the optimal secretion signal. To further validate its utility, we tested αPrePro-driven agonism and confirmed that it supported C5a activation of C5aR, CART peptide activation of GPR68, and TRV056 peptide activation of AGTR1 (Fig. S4g). To estimate the amount of autocrine ligand secretion, we next compared autocrine signaling outputs to dose-response curves generated with exogenously added peptide agonists. Specifically, we performed α-factor titrations of Ste2 and SRIF-14 titrations of SSTR5 and matched the efficacy and agonist concentrations between exogenous stimulation and autocrine activation. Based on this comparison, we estimated that the concentration of secreted autocrine ligand ranged between approximately 0.1 and 1 μM (Fig. S4h).

### PIONEERed luciferase complementation for detecting surface expression

Having established PIONEER, we next sought to demonstrate its utility across different applications. We began by adapting a split luciferase system to detect cell surface expression. This system uses two fragments: LgBiT, an ~18 kDa inactive luciferase, and HiBiT, an 11-amino-acid peptide that binds LgBiT with high affinity to restore activity^34^. As shown in Fig. 2a, we tagged the extracellular N-termini of GPCRs with HiBiT and periplasmically displayed LgBiT using an αPrePro-LgBiT-Ccw14 construct. Surface-localized GPCRs were expected to bring the HiBiT tag into proximity with the Ccw14-displayed LgBiT, reconstituting luciferase activity and producing a detectable signal.

**Fig. 2.**
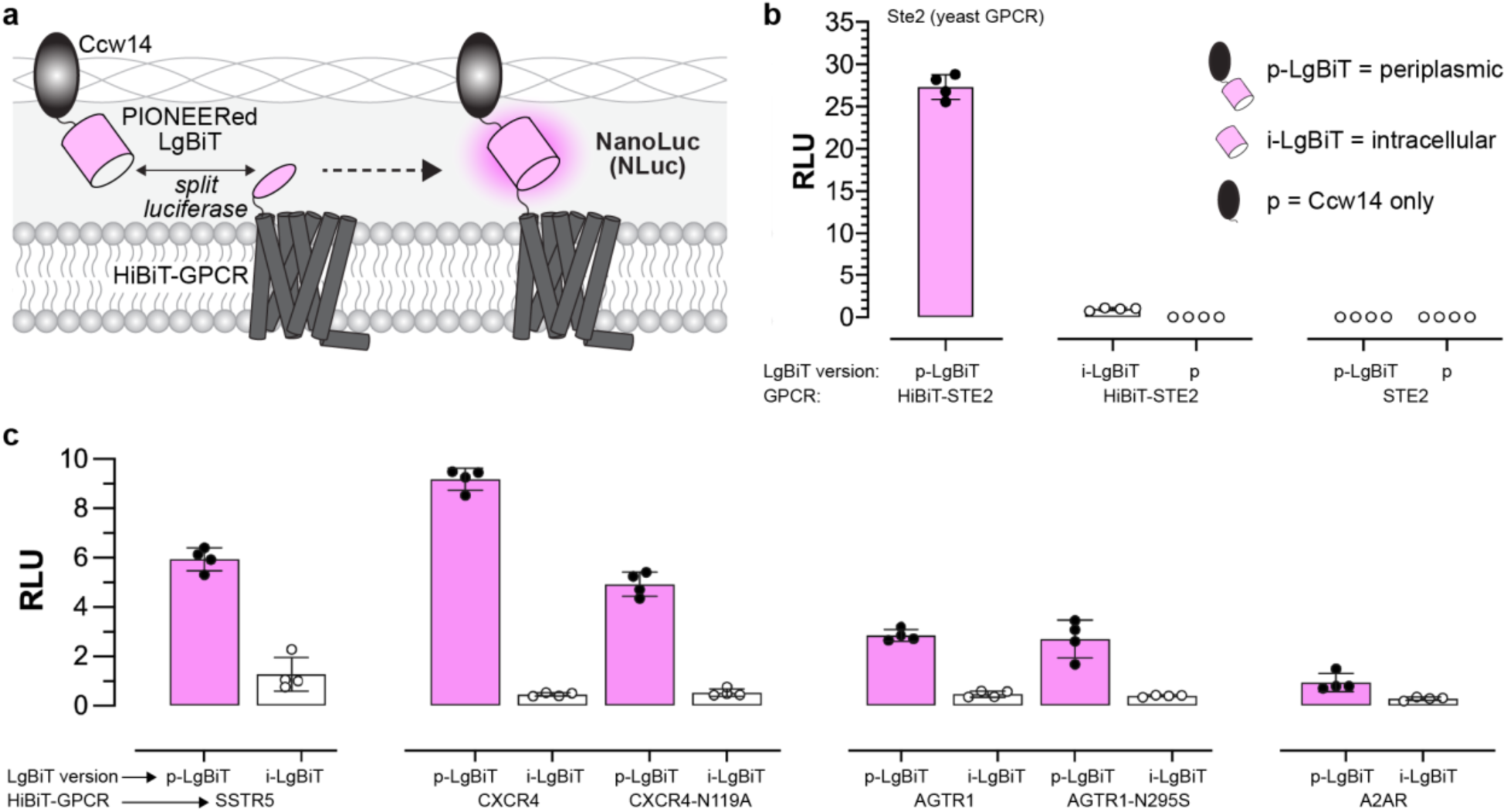
Detection of GPCR surface expression using the PIONEER platform. **a**, Schematic of surface detection using PIONEER. Periplasmically displayed LgBiT enables reconstitution of NanoLuc luciferase and luminescence upon interaction with GPCRs tagged at the N-terminus with HiBiT. **b**, Surface detection of the native yeast GPCR *Ste2* via luminescence reconstitution. **c**, Detection of human GPCRs expressed at the yeast surface using the same approach. Data in **b** and **c** are presented as mean ± s.d. of *n* = 4 biological replicates. Luminescence is reported in relative luminescence units (RLU).

As shown in Figs. 2b and S5, we observed robust luminescence for our control receptor, a HiBiT-tagged version of the yeast GPCR Ste2. Additional controls, including intracellular LgBiT (i-LgBiT) paired with HiBiT-Ste2, periplasmically displayed LgBiT (p-LgBiT) paired with untagged Ste2, and periplasmically displayed Ccw14 alone (p), produced no detectable signal. These results confirmed that luciferase activity is reconstituted only in the full PIONEER configuration when HiBiT binds LgBiT in the periplasmic-like space. Furthermore, these findings establish that LgBiT is a compatible Ccw14 fusion partner.

As shown in Fig. 2c, we evaluated our surface detection method using a panel of human GPCRs that included strong (SSTR5, A2AR, and point mutants CXCR4-N119A and AGTR1-N295S) and weak (wild-type CXCR4 and AGTR1) DCyFIR signalers^21,23,24^. This approach allowed us to directly test whether surface expression correlated with signaling output. Periplasmically displayed LgBiT (p-LgBiT) complemented with all receptors and produced measurable luminescence relative to intracellular LgBiT (i-LgBiT) controls. CXCR4 produced the highest luminescence despite its weak signaling activity, whereas A2AR produced the lowest despite its strong signaling profile^23,24^. These results demonstrate the broad applicability of the assay and reveal that apparent surface expression does not necessarily predict GPCR signaling strength.

### PIONEERed agonists autocrine activate human GPCRs

For our second application, we revisited autocrine agonism of CXCR4 and SSTR5 using the final PIONEER configuration, which combines the αPrePro-ligand-Ccw14 display construct with the DCyFIR system. As shown in Fig. 3a, we employed a panel of ten DCyFIR strains designed to test all receptor-G_α_ coupling combinations. Each strain expresses a human-yeast G_α_ chimera, created by replacing the last five residues of the yeast G_α_ C-terminus with those from a human G_α_ subunit. These chimeras span the G_i/o_ (i, o, t, z), G_q_ (q, 14, 15), G_s_ (s), and G_12/13_ (12, 13) families, enabling human GPCRs to couple efficiently to the yeast signaling pathway. As such, the DCyFIR system supports sustained activation, comprehensive pharmacological studies, and ligand/drug screening, and it incorporates an mTq2 transcriptional reporter as a fluorescent readout.

**Fig. 3.**
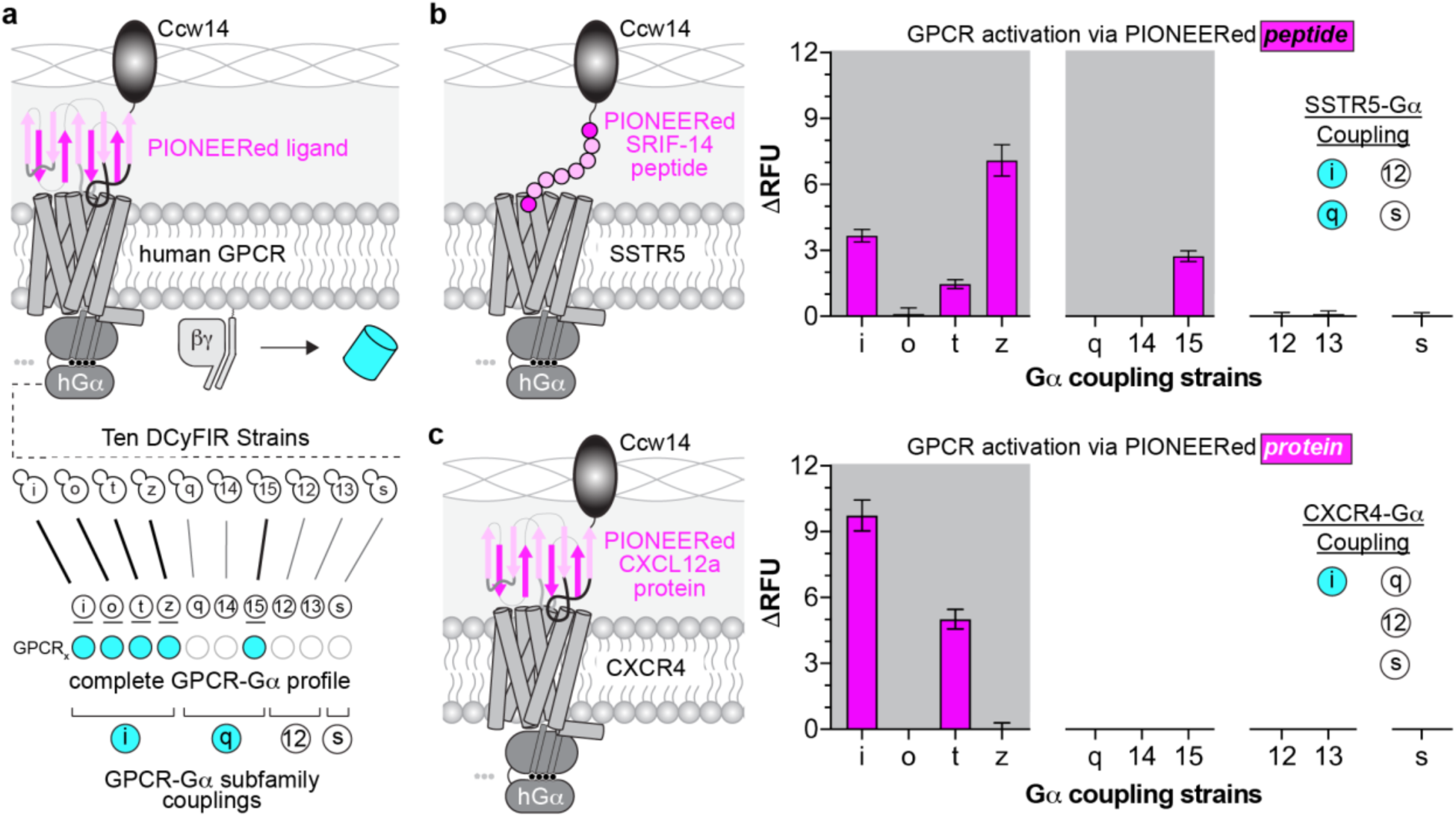
Autocrine activation of GPCRs using the PIONEER platform. **a**, Schematic of autocrine GPCR activation and downstream G protein profiling using the DCyFIR system. Ligand display via PIONEER enables GPCR-G_α_ coupling to be assessed by transcriptional reporter output. **b**, Schematic (left) and functional profiling (right) of autocrine GPCR activation using the PIONEERed peptide ligand, SRF-14. **c**, Schematic (left) and functional profiling (right) using the PIONEERed chemokine ligand, CXCL12a. GPCR activation was measured by mTq2 fluorescence and is shown as ΔRFU relative to the same strain expressing the GPCR and secretion signal only (no encoded ligand). Data represent mean ± s.d. of *n* = 4 biological replicates.

As shown in Figs. 3b-c, SSTR5 signaled through multiple G_i/o_ family members, including G_i_, G_t_, and G_z_, with G_z_ producing the strongest response. It also coupled to the G_q_ family member G_15_. CXCR4 signaled through G_i_ and G_t_, with G_i_ yielding the highest activation. These results demonstrate that the platform is robust, selective, and compatible with both peptide and protein agonists. A key advantage is that genetically encoded ligands can be produced directly in yeast through plasmid transformation or CRISPR integration, eliminating the need for external synthesis or purification. Because the ligand is retained in the periplasm, each receptor is paired with a single ligand, making the system well-suited for high-throughput library screening.

### PIONEERed nanobodies modulate human GPCR signaling

Next, we tested the compatibility of nanobody regulators with the PIONEER system. Nanobodies are single-domain antibody fragments derived from camelid heavy-chain-only antibodies that retain full antigen-binding capacity in a compact and stable format. Their small size and robustness enable efficient production in microbial systems, such as yeast, and access to epitopes often inaccessible to conventional antibodies. They are widely used in GPCR studies to stabilize specific receptor conformations, facilitating structural determination^16^. They also serve as conformational biosensors, enabling the detection and modulation of GPCR activation states in live cells^35^.

We reasoned that the PIONEER system would enable functional characterization of nanobody binding to GPCRs, rather than limiting analysis to binding alone. Conventional yeast surface display presents nanobodies on the outside of the cell wall and relies on purified, bead-bound GPCRs, allowing nanobodies to bind either face of the receptor without functional specificity and providing no signaling readout. In contrast, PIONEER expresses and displays nanobodies in the periplasm, linking nanobody engagement directly to GPCR signaling (Fig. 4a). This configuration enables selective targeting of nanobodies to either the extracellular face (extrabodies) or intracellular face (intrabodies) of the receptor, offering spatial control and functional resolution.

**Fig. 4.**
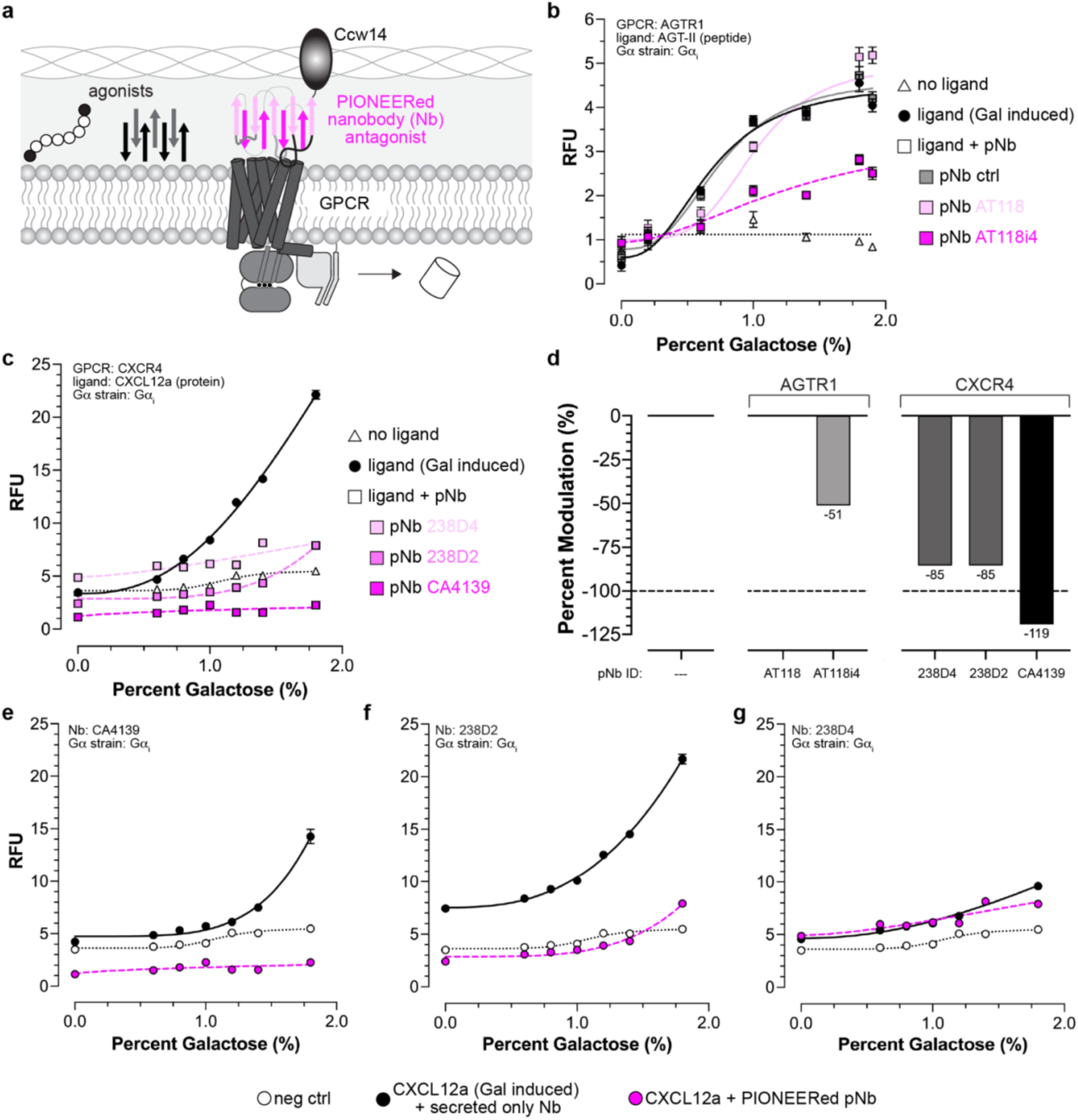
Modulation of GPCR activity using PIONEERed nanobodies. **a**, Schematic illustrating preplasmic nanobody display (pNb) via PIONEER for antagonizing GPCRs that recognize peptide or protein ligands. **b**, Inhibition of AGTR1 signaling by nanobodies displayed using PIONEER. **c**, Inhibition of CXCR4 signaling by PIONEERed nanobodies. GPCRs were activated by secretion of αPrePro-tagged ligands under galactose induction. **d**, Quantification of nanobody-mediated inhibition of AGTR1 and CXCR4 from panels **b** and **c**. **e-g**, Comparison of CXCR4 inhibition by nanobodies with (magenta) and without (black) PIONEER configurations, including CA4139 (**e**), 238D2 (**f**), and 238D4 (**g**). Cells expressing CXCR4 without nanobody constructs served as a negative control. In panels **b** and **c**, nanobody BV025 was used as a periplasmic nanobody (pNb) control. Data represent mean ± s.d. of *n* = 4 biological replicates.

As shown in Figs. 4b-d and S6, PIONEERed extrabodies effectively inhibited AGTR1 and CXCR4 activation. To set up the assay, we placed αPrePro-AGT-II and αPrePro-CXCL12a constructs under a galactose promoter to drive inducible agonist production. By titrating galactose, we generated dose-response curves, modulating the amount of agonist released into the periplasmic space. Antagonist PIONEERed extrabodies were constitutively expressed and localized simultaneously with AGT-II and CXCL12a induction, ensuring receptors were exposed to both agonists and inhibitors. Because ligand release and extrabody engagement occurred in parallel, the assay inherently measured competition for receptor binding, enabling direct quantification of functional inhibition mediated by the PIONEERed nanobodies.

As shown in Fig. 4b, we tested two previously developed extrabodies targeting AGTR1. The first, AT118, is a parent nanobody that binds AGTR1 but does not antagonize its activity, while the second, AT118i4, is a derivative containing two mutations in the CDR1 region that enable competitive antagonism *in vitro* and in an *in vivo* mouse model^36^. Consistent with prior reports, AT118 did not inhibit AGTR1 signaling, although we observed a slight rightward shift in the dose-response curve, suggesting weak inhibition or negative allosteric modulation. As anticipated, AT118i4 robustly inhibited AGTR1 signaling, although it did not completely block receptor activation at higher agonist levels.

As shown in Fig. 4c, we next extended our studies to three extrabodies previously developed for CXCR4. Two of these, 238D2 and 238D4^37^, competitively inhibited CXCR4-mediated signaling and antagonized the chemoattractant activity of CXCL12a. The third, CA4139^38^, was previously reported to enhance CXCR4 expression in *Pichia pastoris*, presumably by acting as a pharmacological chaperone, and in our assay competitively inhibited CXCL12a binding and reduced basal CXCR4 signaling, indicating that it functions as both an inverse agonist and competitive inhibitor. In contrast to AGTR1 inhibition by AT118i4, all three CXCR4 extrabodies maintained inhibition even at higher agonist levels. Together (Fig. 4d), these findings demonstrate the utility of the PIONEER platform for functionally screening and assessing GPCR antagonism by nanobodies.

To complete our analysis of PIONEERed extrabodies, we next tested whether periplasmic display was necessary. As shown in Figs. 4e-g, we examined the three CXCR4 extrabodies (CA4139, 238D2, 238D4) with and without their PIONEER configurations. CA4139 and 238D2 required display to inhibit signaling, whereas 238D4 did not. This suggested that 238D4 might bind stronger than CA4139 and 238D2. However, previous studies reported similar 238D2 and 238D4 EC_50_ values for CXCL12 ligand displacement, CXCR4-G_αqi5_-stimulated inositol phosphate accumulation, and CXCR4-G_αi_-mediated inhibition of cAMP reporter activity^37^. Possible explanations include a slower off-rate for 238D4 or its higher production and secretion in the PIONEER system. Regardless, these findings demonstrate that PIONEER is necessary for capturing nanobody-GPCR interactions across a broad range of kinetic and thermodynamic properties relevant to developing pharmacological tools, biosensors, and preclinical leads.

### Intrabodies modulate GPCR signaling

For our final application, we leveraged PIONEER’s ability to produce nanobodies that can be secreted and displayed as extrabodies or retained inside the cell as intrabodies, which offer several advantages for GPCR studies. Intrabodies can stabilize receptors in specific active or inactive states for high-resolution structural analysis, trap transient signaling intermediates such as GPCR-G protein or GPCR-arrestin complexes, and serve as live-cell biosensors to monitor real-time conformational changes and receptor activity^35,39,40^. To explore these capabilities, we evaluated previously reported intrabodies lacking PIONEER configurations (Fig. 5a), first focusing on AGTR1, for which we had already extensively characterized extrabody counterparts (Fig. 4b).

**Fig. 5.**
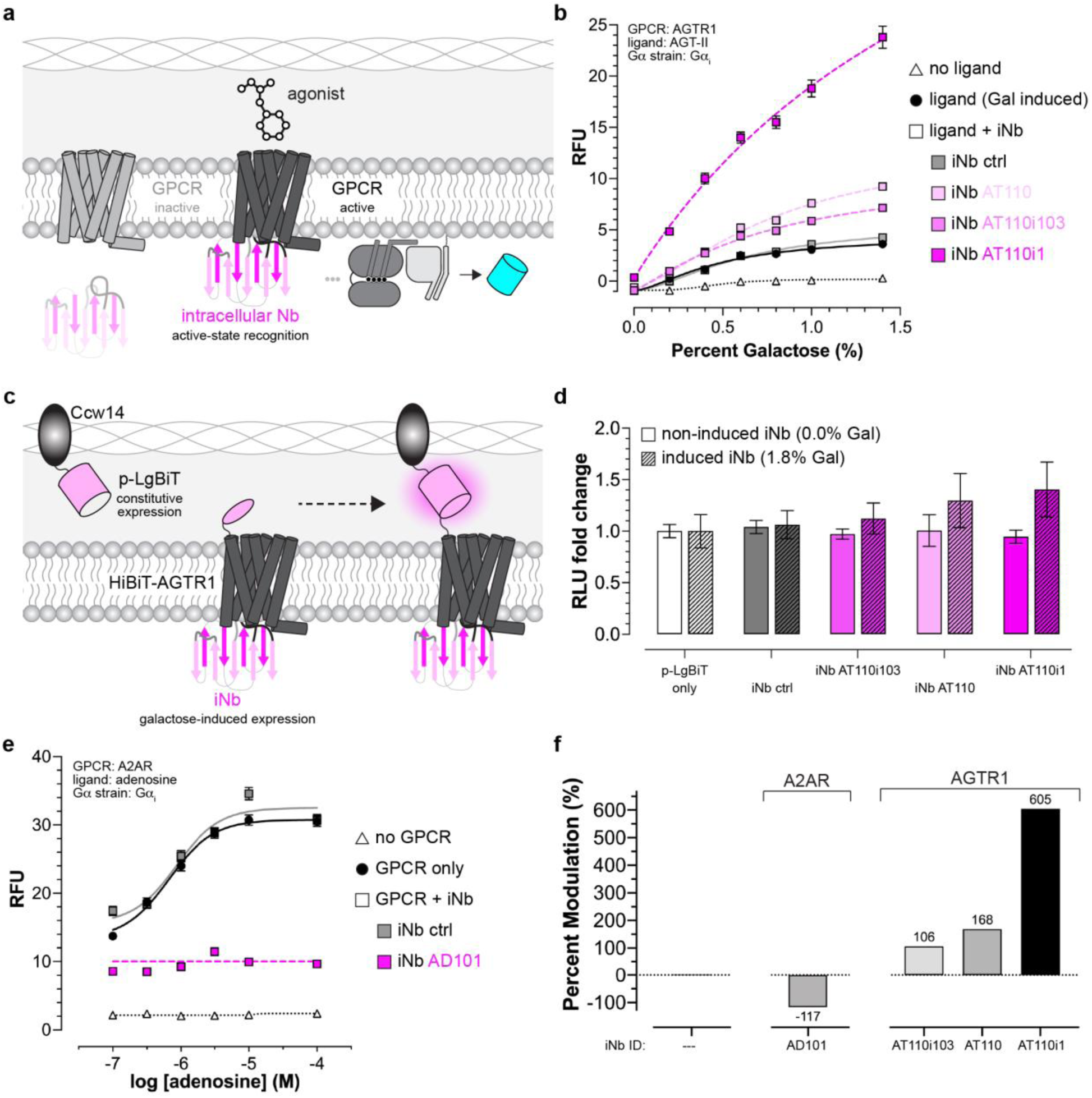
Intracellular nanobody-mediated regulation of GPCR signaling. **a**, Schematic of the platform adaptation to include intracellular nanobody (iNb) expression for modulating GPCR activity. **b**, Intrabodies that enhance AGTR1 signaling upon activation by a secreted αPrePro-AGT-II ligand. **c**, Schematic of PIONEER-based surface detection for monitoring iNb effects on AGTR1 surface expression. **d**, Changes in AGTR1 surface expression upon induction of iNbs used in **b**. **e**, Intrabodies that inhibit A2AR signaling in response to exogenous adenosine. **f**, Quantification of nanobody-mediated modulation of AGTR1 and A2AR signaling from panels **b** and **e**. In panels **b**, **d**, and **e**, nanobody BV025 was used as an iNb control. Data represent mean ± s.d. of *n* = 4 biological replicates.

As shown in Figs. 5a and S7, we evaluated three previously described AGTR1 intrabodies. The parental nanobody, AT110, and its enhanced variant, AT110i1, were developed for structural studies to bind the intracellular pocket of AGTR1 and stabilize its active conformation^41^. A third intrabody from a separate study, AT110i103, was developed using autonomous hypermutation yeast surface display (AHEAD) and exhibited potent nanomolar allosteric modulation^13^. In all cases, intrabody binding was characterized using radioligand allosteric shift assays, but their effects on AGTR1 signaling remained unclear.

To address this, we applied the same approach as in our extrabody studies of AGTR1. We generated dose-response curves by titrating galactose to induce AGT-II agonist expression and its secretion into the periplasmic space, with intrabodies constitutively expressed and co-localized during induction. Based on their binding to the active receptor conformation, we expected the intrabodies would inhibit signaling by competing for AGTR1’s G_α_ binding site and preventing G protein engagement. However, all three intrabodies unexpectedly enhanced pathway output in response to AGT-II.

As shown in Fig. 5b, intrabodies AT110 and AT110i103 modestly enhanced AGTR1 signaling, approximately doubling output at peak AGT-II levels, while AT110i1 had a much stronger effect, increasing signaling more than tenfold. None of the intrabodies increased basal signaling, indicating they do not disrupt the G protein heterotrimer to release G_β_. Instead, they likely function as pharmacological chaperones, similar to the CXCR4 extrabody CA4139, by increasing the amount of receptor available for activation. If so, stronger chaperoning should correlate with greater signaling output, supporting the use of this platform to efficiently identify active-state stabilizing intrabodies for various GPCRs.

To test this, we used our PIONEER-based surface detection system to monitor changes in surface AGTR1 when iNb expression was induced (Fig. 5c). As expected, we observed an increase in AGTR1 surface levels when either AT110, AT110i103, or AT110i1 were expressed (Fig. 5d), whereas no changes were detected with our control iNb. Furthermore, the relative changes in AGTR1 levels correlated with our signaling analyses from Fig. 5b, with AT110i1 invoking the greatest effects. Together, these data supported our chaperoning hypothesis and emphasize the utility of this platform for identifying GPCR-stabilizing intrabodies.

Lastly, as shown by the adenosine dose-response curves in Figs. 5e and S7a, we tested an A2AR intrabody, AD101. Like the AGTR1-directed nanobodies, AD101 recognizes the agonist-bound active state^16^. However, unlike its AGTR1 counterparts, AD101 acted both as an inverse agonist and completely inhibited signaling, likely by blocking G protein recruitment. This suggests that AD101 does not increase receptor abundance by chaperoning. These results demonstrate that the platform can distinguish active-state intrabodies based on both their functional effects and chaperone ability. Intrabodies that do not alter receptor abundance may be better suited for *in vivo* biosensing, while chaperoning intrabodies could enhance dynamic range in engineered strains or support identification of high-affinity binders for structural studies. A summary of both intrabody-receptor results is shown in Fig. 5f.

## Discussion

### Advancing periplasmic display for synthetic biology

The results presented here establish PIONEER as a versatile platform for functional screening of genetically encoded ligands and proteins using yeast periplasmic display, a conceptually rich but underexplored tool in synthetic biology. Unlike yeast surface display, which presents proteins on the outer cell wall for endpoint assays, periplasmic display localizes them to a confined, membrane-proximal space suited for functional studies. This approach is especially useful for peptides and proteins that are difficult or expensive to produce in heterologous systems and offers a scalable platform for studying targeted and naïve libraries of protein-protein interactions across diverse biotechnological applications.

Importantly, the effectiveness of PIONEERed constructs does not depend on their ability to span the periplasmic space. Ccw14-mediated cell wall binding is dynamic, with on/off kinetics that allow ligand diffusion within the compartment while preventing cargo escape. We speculate that signaling is primarily mediated by freely diffusing constructs rather than those statically tethered to the cell wall, enabling more flexible and repeated engagement with membrane-embedded targets. Building on these insights, we are exploring periplasmic display scaffolds compatible with ligands that engage their targets via C-terminal interactions to further expand the functional utility of the PIONEER platform.

### GPCR-specific advances and future applications

Our results also demonstrate several advances for GPCR research. First, we show that PIONEERed agonists can activate human GPCRs with defined G_α_-coupling preferences, enabling comprehensive pharmacological profiling. Second, we demonstrate that intrabodies and PIONEERed extrabodies provide spatial precision for modulating GPCR activity through stabilization, inhibition, and chaperoning. Third, we establish that the system can distinguish functional classes of nanobodies, including inverse agonists, competitive antagonists, and active-state stabilizers. These capabilities make PIONEER broadly applicable to drug discovery, biosensor development, and structural biology, where identifying conformationally selective ligands remains a significant challenge. The ability to assess peptide and protein ligand activity through signaling, rather than binding alone, marks a key advance toward function-first screening.

While the present study establishes feasibility and core functionality, we acknowledge that the potential for large-scale, fluorescence-based discovery has not yet been explicitly demonstrated. Nonetheless, our prior work with the DCyFIR platform, on which PIONEER builds, has shown that fluorescence-activated cell sorting (FACS) enables robust, specific, and quantitative enrichment of functional variants in pooled library formats. These precedents support the broader scalability of PIONEER for high-throughput screening, while underscoring that the current work focuses primarily on validating the foundational framework.

### Combining AI and PIONEER

AI models are increasingly capable of generating peptides, nanobodies, and protein variants predicted to modulate biological function, yet most of these predictions remain untested in living systems. Without experimental grounding, even high-confidence models risk producing sequences that are biophysically plausible but biologically inert. PIONEER addresses this challenge by providing a genetically encoded, scalable platform that links designed sequences to functional output in a cellular context. By enabling the direct testing of AI-generated molecules for activity, inhibition, or stabilization, PIONEER delivers the functional resolution needed to evaluate and refine computational designs.

Although this study focuses on GPCRs, the PIONEER framework is agnostic to target class and broadly applicable to diverse membrane proteins, interaction partners, and signaling pathways. For example, a variety of mammalian receptor tyrosine kinases (RTKs) retain function when expressed in yeast^42^ and have been studied using the first-generation Flo42-based autocrine design^43^. As such, PIONEER can be readily adapted to these systems with minimal modification or integrated with GPCR-centric platforms such as DCyFIR to investigate GPCR–RTK crosstalk.

More generally, PIONEER provides a conceptual and experimental foundation for studying interactions between any genetically encodable proteins. While large-scale implementation will require additional optimization and benchmarking, the framework is compatible with high-throughput readouts such as FACS and other pooled selection strategies. Two examples include using the platform to explore and optimize split reporter systems or to design new enzyme–substrate pairs. In each case, productive complementation or enzymatic activity could, in principle, be resolved in high-throughput formats, building on the robust sorting and selection strategies established in our previous work. As AI-driven design continues to expand into more complex biological systems, frameworks like PIONEER that assess molecular function rather than only binding or structure will be essential.

## Experimental Procedures

### Reagents

All media, buffers, and solutions used throughout this work are described in Supplementary Table 1.

### Cell culture

NEB® 5-alpha Competent *Escherichia coli* (*E. coli*), a derivative of the DH5α strain, were used for all cloning experiments and general plasmid propagation.

All yeast strains used throughout this study are derivatives of BY4741 (MATa *his3Δ1 leu2Δ0 met15Δ0 ura3Δ0*) and are listed in Supplementary Table 2. Specifically, the DCyFIR DI3Δfig1Δ::mTq2 P1 X-2 model strain (BY4741 *far1Δ0 sst2Δ0 ste2Δ0 fig1Δ0::mTq2 X-2:P_TEF1a_-UnTS-T_CYC1b_*) or derivatives were used and have been described previously ^24^. Prior to transformation or other experimentation, yeast cells were grown at 30°C on Yeast extract Peptone Dextrose (YPD) agar medium. Select colonies were propagated at 30°C, static or shaking at 200 rpm, in liquid cultures of YPD or pH-adjusted Synthetic Complete Dextrose low fluorescence medium (SCD Screening Medium; pH 7.0) as specified. Transformations were performed using a standard lithium acetate protocol described previously ^21^. Transformants were initially selected for at 30°C for three days on SCD dropout agar medium. Select transformants were then propagated at 30°C, static or shaking at 200 rpm, in liquid cultures of SCD dropout medium. For all absorbance (*A*_600_) normalization steps, a Biomek NX^P^ liquid-handling robot was used.

### Display protein informatics

Sequences for native yeast display proteins (see Supplementary Fig. 2a) were obtained from the *S. cerevisiae* strain S288C, the parental BY4741 strain, reference genome sequence R64-2-1 via the *Saccharomyces* Genome Database (SGD)^44^. Protein sequences were analyzed using the SignalIP 5.0 ^45^ and NetGPI 1.1 ^46^ software to identify native secretion signals and glycosylphosphatidylinositol (GPI) anchor features present, respectively. An engineered construct with a known secretion signal and GPI anchor (Signal-NbBV025-649stalk) ^16^, as well as the native Gγ protein Ste18, a well-studied membrane-associated protein with no secretion sequence or GPI, were used for initial software validation.

### Cloning and molecular biology

All plasmids used and constructed throughout this study are listed in Supplementary Table 3 and were validated via Sanger sequencing (Eurofins Genomics). All GPCR constructs were generated in a pYEplac195 vector using NEBuilder® HiFi DNA Assembly Master Mix (New England Biolabs). Genes were sourced from the BY4741 genome (STE2) or the PRESTO-TANGO library (WT human GPCRs) ^47^. CXCR4-N119A and AGTR1-N295S mutants were first created in the PRESTO-TANGO vector via site-directed mutagenesis then cloned into pYEplac195 via HiFi DNA Assembly. These constructs were used for all GPCR assays except for Figure 2c and Fig. S5, as noted, as no signaling was detectable with WT versions.

Plasmids used for initial testing of display proteins were constructed by amplifying the respective genes from the BY4741 genome, starting immediately downstream of the secretion signal cleavage site (see Supplementary Fig. 2a). The one exception is the Flo42 construct, which contained only the C-terminal 42 amino acids of Flo1 and was previously synthesized as a gBlock by Integrated DNA Technologies (IDT) ^21^. Amplicons were cloned into pYEplac181 αPre via HiFi DNA Assembly as αPre-*X*, where *X* is the display candidate. A yeast codon optimized mTurquoise2 (Addgene #86424), mTq2 ^24,48^, was then inserted between αPre and the display protein gene in similar fashion for mTq2-based analyses (see Figs. 1c-d and Supplementary Fig. 2b-c). For initial GPCR autocrine activation (Figs. 1d-e and Supplementary Fig. 3), SRIF-14 and CXC12a were cloned into pYEplac181 αPre-*X*, where *X* is the display candidate, via inverse PCR ^25^ or HiFi DNA Assembly, respectively, as αPre-*Y*-*X*, where *Y* is the genetically encodable ligand. The CXCL12a gene was codon optimized for *S. cerevisiae* (IDT) and sourced as a gBlock.

For mTq2-based secretion signal (ss) testing, six unique signals were selected for comparison with aPre (see Fig. 1f, Supplementary Fig. 4a and Supplementary Table 3). Complete secretion signals were first generated via Assembly PCR using 2-4 overlapping oligonucleotides 40-100 bp long. Once assembled, amplicons were directly cloned into pYEplac181 as described above, followed by cloning of mTq2 downstream of the secretion signal. For ligand-based secretion signal testing (see Fig. 1f and Supplementary Figs. 4d-g), pYEplac181 *ss*-*Y* constructs were generated via inverse PCR (α-factor, SRIF-14, AGT-II, CART (42-89)_(9-28), and TRV056) or HiFi DNA Assembly (CXCL12a and C5a), where *Y* is the genetically encodable ligand. The C5a gene was codon optimized for *S. cerevisiae* and sourced as a gBlock.

For PIONEER-based detection of surface GPCRs, pYEplac195 *GPCR* were N-terminally tagged with the HiBiT peptide followed by a GGSG linker via inverse PCR. Constructs for expressing cytosolic or PIONEERed LgBiT were created by cloning LgBiT, sourced from pBiT1.1C-LgBiT (gift from Nevin Lambert), into pYEplac181 or pYEplac181 αPrePro-Ccw14 as αPrePro-LgBiT-Ccw14 via HiFi DNA Assembly.

Co-expression of secreted ligands and nanobodies required designing a vector with two expression cassettes; one for constitutive expression (PIONEERed or intracellular nanobody) and one inducible (secreted ligand). Starting with pYEplac181 (constitutive *TEF1* promoter and *CYC1* terminator), an additional cassette containing the inducible *GAL1* promoter and *ADH1* terminator were inserted upstream of *TEF1* in reverse orientation via HiFi DNA Assembly, similar to a construct described previously ^49^. We refer to this construct as pYEplac181M2. *GAL1* and *ADH1* sequences were sourced from pYDS649 ^16^. Secreted ligands and nanobodies (PIONEERed, αPrePro-*nanobody*-Ccw14; intracellular, *nanobody*) were then cloned into the *P_GAL1_*-*T_ADH1_* and *P_TEF1_*-*T_CYC1_* expression cassettes, respectively, via HiFi DNA Assembly. For monitoring intracellular nanobody effects on GPCR surface expression (Figs. 5c-d), nanobodies and αPrePro-LgBiT-Ccw14 were cloned into the *P_GAL1_-T_ADH1_* and *P_TEF1_-T_CYC1_* expression cassettes, respectively. In instances where GPCR activation was achieved via exogenous ligand application (A2AR with adenosine; see Figs. 5e-f, and Fig. S7a), nanobodies were expressed from the parental pYEplac181 vector. All nanobody sequences were codon optimized for *S. cerevisiae* and sourced as gBlocks.

### Data acquisition

A CLARIOstar multimode microplate reader (BMG LabTech) was used for collecting all microplate-based absorbance at 600 nm (*A*_600_), temperature, fluorescence (RFU), and luminescence (RLU) data with the following instrument settings: *A*_600_ (excitation 600 nm; 22 flashes/well), mTq2-based fluorescence (excitation 430/10 nm, dichroic LP 458 nm, emission 482/16 nm; 10 flashes/well; bottom read), NanoLuc luminescence (emission 482/16 nm; top read).

### Yeast growth/fitness assay

Yeast cells transformed with pYEplac181 αPre-mTq2-*X*, where *X* is the display candidate, were picked from SCD-LEU agar medium into 500 uL SCD-LEU medium in sterile 96-well deep-well blocks (Greiner Bio-One), covered with porous film (Diversified Biotech), shaken on a MixMate microplate shaker (1200 rpm for 30 sec) (Eppendorf), and placed at 30°C. After 18 hrs, cultures were used to prepare new 500-uL cultures normalized to *A*_600_ 0.2 and allowed to grow at 30°C for 3 hrs. Finally, 50-uL cultures normalized to *A*_600_ 0.1 were prepared in a black 384-well clear-bottom plate (Greiner Bio-One) and placed in a microplate reader programmed for 30°C incubation. Automated temperature and *A*_600_ data were collected every hr for 15 hrs. Data were plotted in GraphPad Prism as time versus *A*_600_ and fit using a standard logistic growth equation.

### Confocal microscopy

In preparation, one yeast colony was picked from SCD dropout agar medium into 5 mL SCD dropout medium and grown overnight at 30°C shaking (200 rpm). Cells were then washed with 5 mL SCD Screening Medium and resuspended in 400 uL, with cell pelleting (3000 g for 3 min) and harvesting prefacing each wash/resuspension step. 2 uL cell suspension was then added to a 75 x 25 x 1-mm microscope slide and covered with a 22 x 22-mm glass coverslip (no. 1.5). Cells were imaged using an LSM800 confocal microscope (Zeiss) at 63x magnification using excitation lasers of 405 nm and processed using Zeiss’s Zen software.

### Display and secretion output and efficiency assays

For determining display protein output and efficiency, yeast strains (DI3Δfig1Δ::mTq2 P1 X-2) transformed with pYEplac181 αPre-mTq2-*X*, where *X* is the display candidate, were picked from SCD-LEU agar medium into 500 uL SCD-LEU medium in sterile 96-well deep-well blocks, covered with porous film, shaken (1200 rpm for 30 sec), and grown at 30°C. After 16 hrs, cultures were used to prepare new 250-uL cultures in sterile clear 96-well plates (CytoOne) normalized to *A*_600_ 0.1 in SCD Screening Medium. Plates were covered with porous film and grown at 30°C for 21 hrs. After this time, 50 uL resuspended cells were transferred to a black 384-well clear-bottom plate. The remaining cultures were centrifuged (3000 g for 5 min) and 50 uL supernatant transferred to empty wells of the same 384-well plate. This plate was then centrifuged briefly, shaken (2000 rpm for 30 sec), and *A*_600_ and fluorescence (gain 850) were measured to determine the degree of mTq2 secreted (supernatant) relative to total mTq2 output (cell suspension). Display efficiency was calculated using Equation 1, where F_s_ is the relative fluorescence of the supernatant for αPre-mTq2-*X* (F_sX_) or empty vector (F_s0_), and F_t_ is the total relative fluorescence of αPre-mTq2-*X* (F_tX_) or empty vector (F_t0_), where *X* is the display candidate.

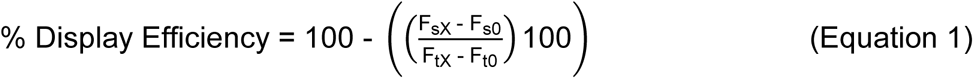

For determining output and efficiency of secretion signals (*ss*), pYEplac181 *ss*-mTq2 constructs were transformed, cells prepared, experiments performed, and data collected as described above. Total fluorescence of the cell suspensions was reported as total protein output, whereas secretion efficiency was calculated using Equation 2. F_sX_ and F_tX_ are the relative fluorescence of the supernatant or total cell suspension of pYEplac181 *ss*-mTq2 transformants, respectively.

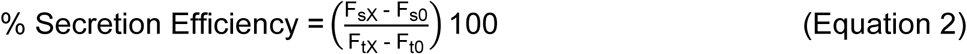

### PIONEER-based split luciferase assay for surface receptor detection

Yeast cells co-transformed with pYEplac181 αPrePro-LgBiT-Ccw14 (p-LgBiT) and pYEplac195 HiBiT-*X*, where *X* is the GPCR, were picked from SCD-URA-LEU agar medium into 1 mL SCD-URA-LEU medium in sterile 96-well deep-well blocks, covered with porous film, shaken (1200 rpm for 30 sec), and placed at 30°C. Co-transformations of HiBiT-GPCRs with intracellular LgBiT (pYEplac181 i-LgBiT) or PIONEER only (pYEplac181 p) and non-tagged GPCRs (pYEplac195 *GPCR*) with pYEplac181 p or p-LgBiT were included as controls (see Figs. 2b-c and Fig. S5). After 21 hrs, cultures were used to prepare new 200-uL cultures normalized to *A*_600_ 0.2 in SCD Screening Medium in sterile clear 96-well plates, covered with porous film, and placed at 30°C. After 15 hrs, cultures were used to prepare 80-uL cultures normalized to *A*_600_ 1.0 in a black, black-bottom 384-well plate (Greiner Bio-One) with furimazine (1:1000). Plate was shaken (2000 rpm for 30 sec) and luminescence measurements (gain 3000) were performed in a microplate reader. Data were plotted as relative luminescence units (RLU) and analyzed in GraphPad Prism.

For monitoring changes in AGTR1 surface levels in the presence of intracellular nanobodies (Figs. 5c-d), yeast cells co-transformed with pYEplac195 HiBiT-AGTR1 and a pYEplac181M2-derived construct dually expressing αPrePro-LgBiT-Ccw14 (*P_TEF1_*-αPrePro-LgBiT-Ccw14*-T_CYC1_*) and a nanobody under galactose regulation (*P_GAL1_*-*nanobody-T_ADH1_*) were prepared in SCD-URA-LEU medium in similar fashion. However, after the initial 21-hr growth, cultures were used to prepare new 1.5-mL cultures in low-glucose SCD-URA-LEU medium (1.9% galactose, 0.1% glucose) in sterile 96-well deep-well blocks normalized to *A*_600_ 0.2. Blocks were covered with porous film, shaken, and placed at 30°C. After 21 hrs, low-glucose cultures were used to prepare new 200-uL cultures normalized to *A*_600_ 0.2 in sterile clear 96-well plates in either SCD Screening Medium without galactose or 1.8% galactose, maintaining a total sugar content of 2.0%. The plate was covered with porous film and placed at 30°C for 15 hrs. Finally, cultures were used to prepare 80-uL cultures normalized to *A*_600_ 1.0 in a white, white-bottom 384-well plate (Greiner Bio-One) with furimazine (1:1000). Plate was shaken (2000 rpm for 30 sec) and luminescence measurements (gain 3000) were performed in a microplate reader. Data were plotted as relative luminescence units (RLU) and analyzed in GraphPad Prism. Fold change in luminescence was calculated relative to p-LgBiT only (no nanobody) in the matched sugar-specific medium.

### GPCR activation assay – exogenous ligand application

For achieving GPCR activation via exogenous ligand application, receptor expression constructs (pYEplac195 *GPCR*) were transformed into the DI3Δfig1Δ::mTq2 P1 X-2 yeast strain (pYEplac195 Ste2) or derived DCyFIR strains (Supplementary Table 2). Resulting transformants were picked from SCD-URA agar medium into 1 mL SCD-URA medium in sterile 96-well deep-well blocks, covered with porous film, shaken (1200 rpm for 30 sec), and placed at 30°C.

After 18-21 hrs, cultures were used to prepare new 72-uL duplicate cultures in a sterile black 384-well clear-bottom plate normalized to *A*_600_ 0.11 in SCD Screening Medium. Following this, 8 uL of 10x drug or vehicle (H_2_O) were added per well. The plate was briefly centrifuged, shaken (2000 rpm for 30 sec), and placed in a microplate reader programmed for 30°C incubation. Automated temperature, *A*_600_, and mTq2 fluorescence (gain 1200) data were collected every hr for 15 hrs.

Data were plotted in GraphPad Prism as fluorescence versus *A*_600_ and fit via linear regression to generate slope and intercept values for each treatment condition, which were used to extrapolate fluorescence to a standardized *A*_600_ of 1.0 and reported as relative fluorescence units (RFU). The resulting RFU data were then plotted as a function of log[ligand] and fit using the four-parameter pharmacological function log(agonist) versus response. Where applicable, error bars represent the standard deviation of the fitted slopes.

### Autocrine GPCR activation assay – secreted and PIONEERed ligands

For initially achieving autocrine GPCR activation, GPCR expression constructs were co-transformed with constructs for ligand secretion (pYEplac181 *ss-ligand*) or secretion and periplasmic display (pYEplac181 *ss-ligand-display*) into the DI3Δfig1Δ::mTq2 P1 X-2 yeast strain (pYEplac195 Ste2) or derived DCyFIR strains. Resulting transformants were picked from SCD-URA-LEU agar medium into 1 mL SCD-URA-LEU medium in sterile 96-well deep-well blocks, covered with porous film, shaken (1200 rpm for 30 sec), and placed at 30°C. After 18-21 hrs, cultures were used to prepare new 80-uL cultures in a sterile black 384-well clear-bottom plate normalized to *A*_600_ 0.1 in SCD Screening Medium and placed in a microplate reader programmed for 30°C incubation. Automated data collection and analyses were performed as described above.

### Nanobody-based modulation of GPCR signaling

For testing GPCR functional modulation by both extracellular-and intracellular-binding nanobodies, GPCR expression constructs were typically co-transformed with pYEplac181M2-derived constructs dually expressing a defined nanobody (*P_TEF1_*-αPrePro-*nanobody*-Ccw14*-T_CYC1_* or *P_TEF1_*-*nanobody-T_CYC1_*) and a secreted ligand under galactose regulation (*P_GAL1_*-αPrePro-*ligand-T_ADH1_*) into specified DCyFIR strains. For instances where ligands were added exogenously, nanobodies were expressed from pYEplac181M2 constructs without encoded ligands (*P_GAL1_-T_ADH1_*). Resulting transformants were picked from SCD-URA-LEU agar medium into 1 mL SCD-URA-LEU medium in sterile 96-well deep-well blocks, covered with film and shaken, and placed at 30°C for 18-21 hrs. Cultures were then used to prepare new 1.5-mL cultures in low-glucose SCD-URA-LEU medium (1.9% galactose, 0.1% glucose) in sterile 96-well deep-well blocks normalized to *A*_600_ 0.2. Blocks were covered with porous film, shaken, and placed at 30°C.

After 18-21 hrs, low-glucose cultures were used to prepare new 80-uL cultures in a sterile black 384-well clear-bottom plate normalized to *A*_600_ 0.1 in one of eight SCD Screening Medium solutions with varying galactose/glucose ratios (see Supplementary Table 1), maintaining a total sugar content of 2.0%. The plate was then placed in a microplate reader programmed to 30°C incubation and automated temperature, *A*_600_, and mTq2 fluorescence (gain 1200) data were collected every hr for 15 hrs.

Data were plotted in GraphPad Prism as fluorescence versus *A*_600_ and fit via linear regression to generate slope and intercept values for each treatment condition, i.e., galactose/glucose ratio (for secreted ligands) or ligand concentration (exogenous ligand applications), which were used to extrapolate fluorescence to a standardized *A*_600_ of 1.0 and reported as RFU. The resulting RFU data were then plotted as either a function of galactose percentage and fit using the four-parameter pharmacological function agonist versus response (secreted ligands), or as a function of log[ligand] and fit using the pharmacological function log(agonist) versus response (exogenous ligand applications). Where applicable, error bars represent the standard deviation of the fitted slopes.

## Supporting information

List of recipes, plasmids, and strains

## Data Availability

All of the data for this study are available in the manuscript and its supporting information.

## Supporting Information

This article contains supporting information.

## Acknowledgements

We thank Nicholas J. Kapolka, Geoffrey J. Taghon, and Sam A. Taylor for their critical input throughout the project. We also thank Aashish Manglik, Andrew Kruse, Aaron Ring, and Chang Liu for insightful early discussions during platform development.

D.G.I. and J.B.R. are inventors on a pending U.S. utility patent application (filed November 29, 2023) related to the PIONEER platform. The other authors declare no competing interests.

## Author Contributions

J.B.R. and D.G.I. conceived the study and co-wrote the manuscript. J.B.R. designed and performed all experiments with the assistance of K.L.

---

**Fig. S1.**
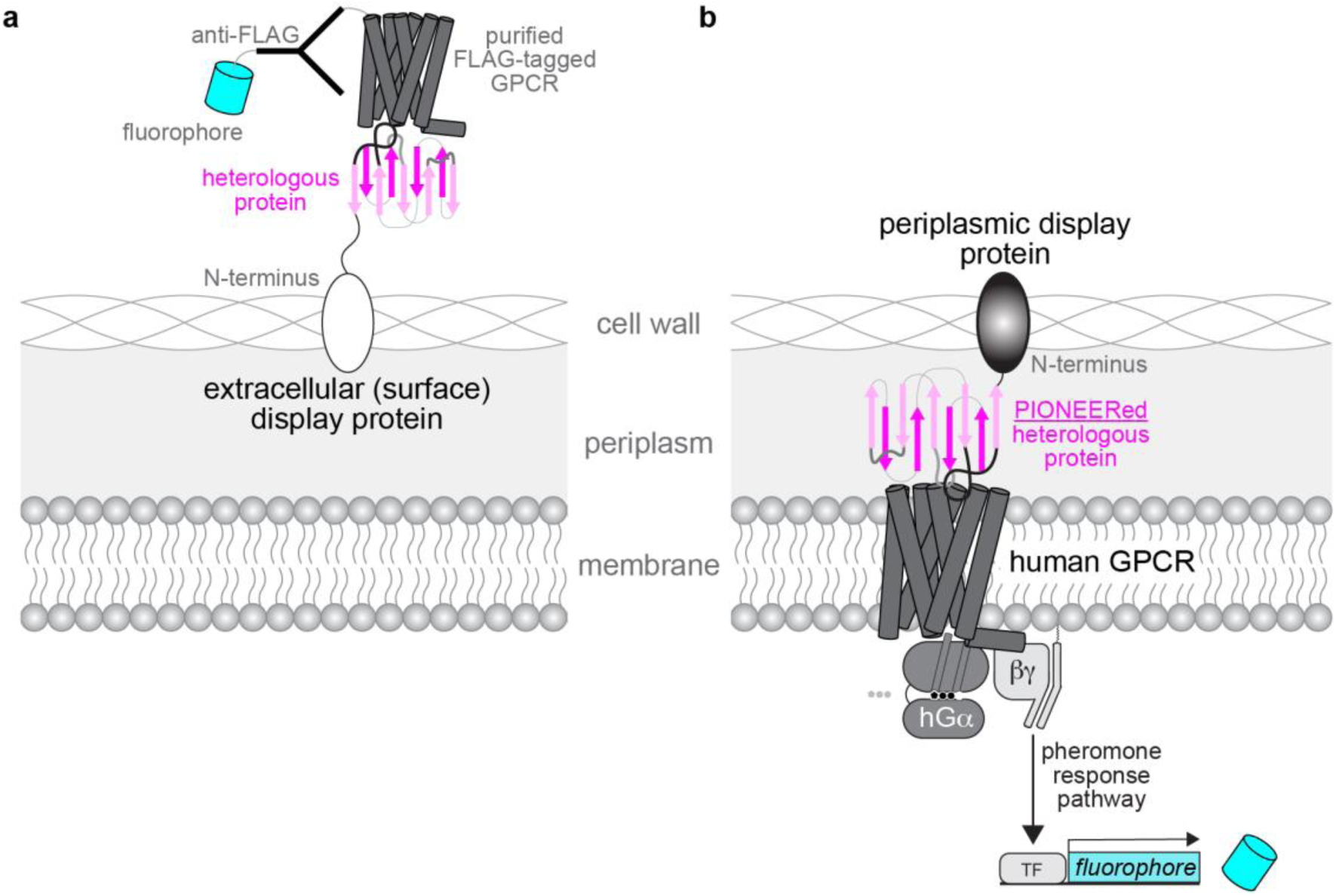
Yeast surface versus PIONEER-based protein display. **a**, Schematic of conventional yeast display systems, where heterologous proteins are presented on the exterior of the cell wall. **b**, Schematic of the PIONEER system, which displays proteins in the periplasmic space between the plasma membrane and cell wall. Pairing PIONEER with functional yeast signaling platforms, such as DCyFIR, enables detection of physiologically relevant protein-receptor interactions.

**Fig. S2.**
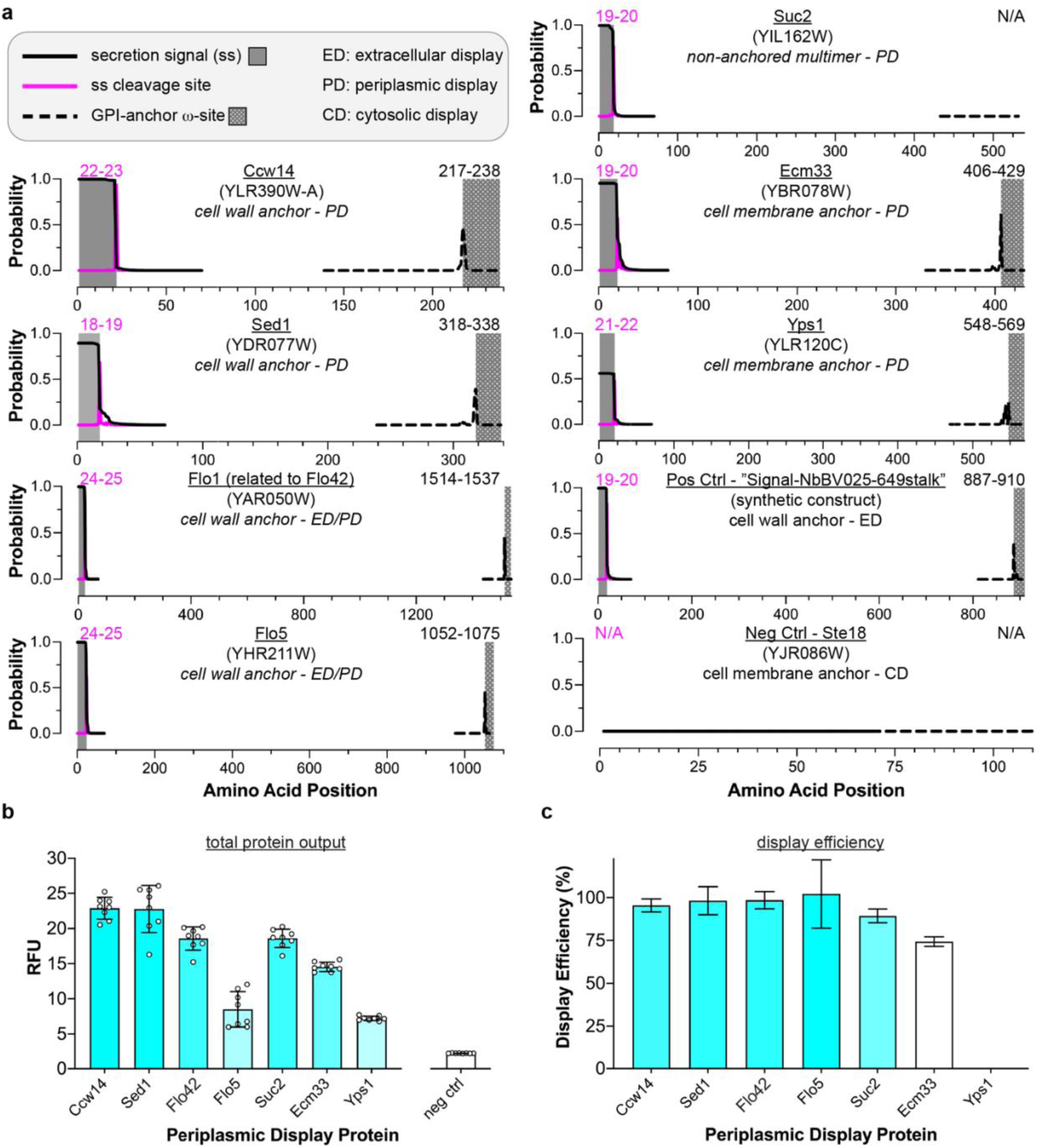
Characterization of periplasmic display proteins for PIONEER. **a**, Sequence analysis of candidate display proteins to identify native secretion signals (black line with grey shading), predicted cleavage sites (magenta line), and glycosylphosphatidylinositol (GPI) anchor motifs (black dashed line with grey mesh shading). Amino acid positions corresponding to cleavage and GPI anchor sites are indicated. Signal-NbBV025-649stalk and *Ste18* were included as positive (synthetic design with known secretion signal and GPI anchor) and negative (membrane-bound, non-secreted) controls, respectively. **b**, Quantification of mTq2 fluorescence from tagged display proteins to assess expression levels. “neg ctrl” indicates cells transformed with empty vector. Data are presented as mean ± s.d. of *n* = 8 biological replicates. **c**, Display efficiency calculated as the percentage of mTq2 fluorescence retained within the cell relative to total mTq2 (intracellular plus secreted). Data are presented as mean ± s.e.m. of *n* = 8 biological replicates. Data from **b** and **c** were used for protein performance scoring in Fig. 1d.

**Fig. S3.**
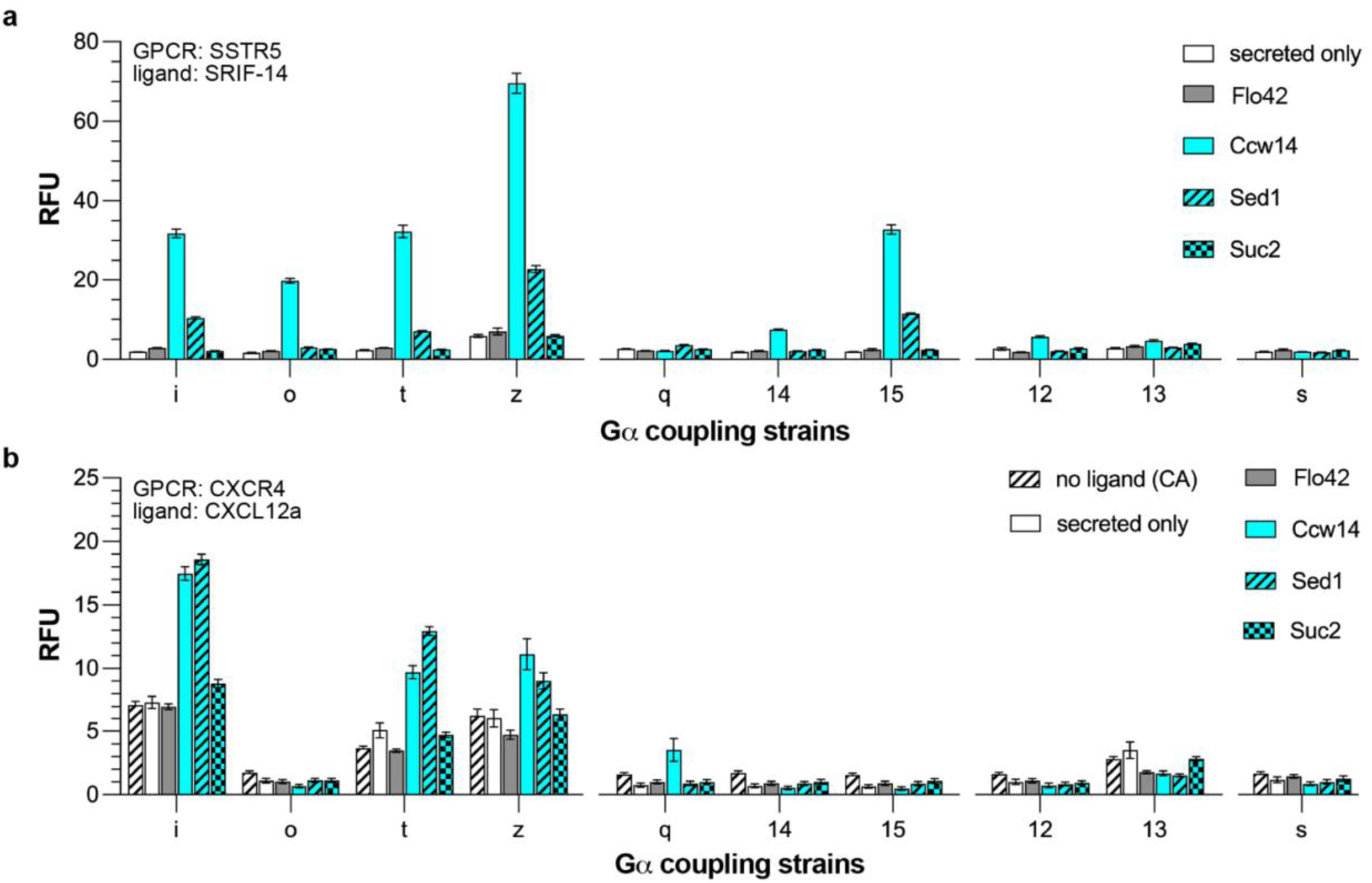
Autocrine GPCR-G_αi_ activation profiles with displayed ligands. **a**, Autocrine activation of SSTR5 by secretion or combined secretion and periplasmic display of its cognate ligand fused to Flo42, Ccw14, Sed1, or Suc2. **b**, Autocrine activation of CXCR4 under the same conditions. Data represent mean ± s.d. of *n* = 4 biological replicates. Experiments were performed using DCyFIR Gαi strains, and results contributed to display protein scoring shown in Fig. 1d-e.

**Fig. S4.**
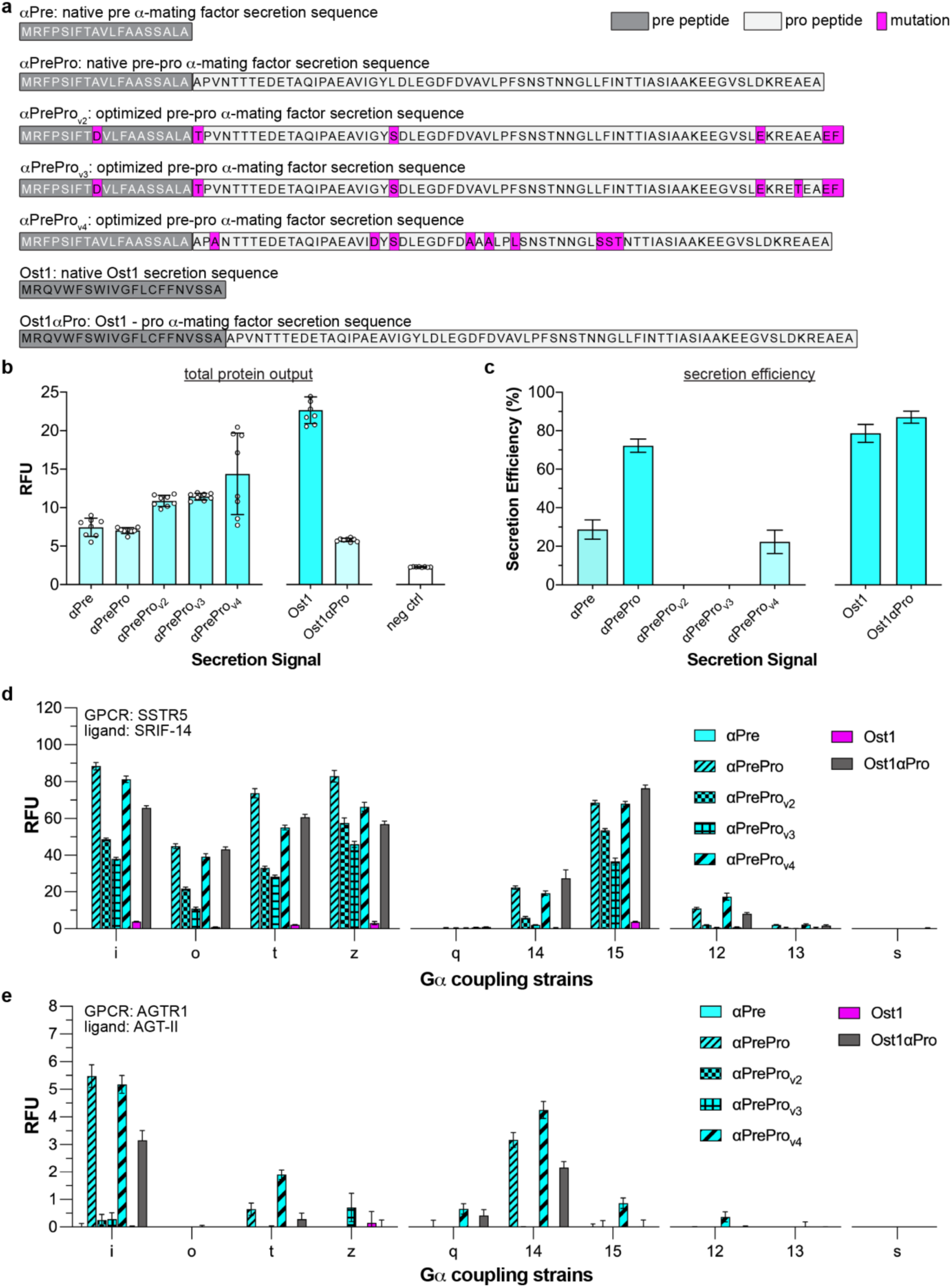
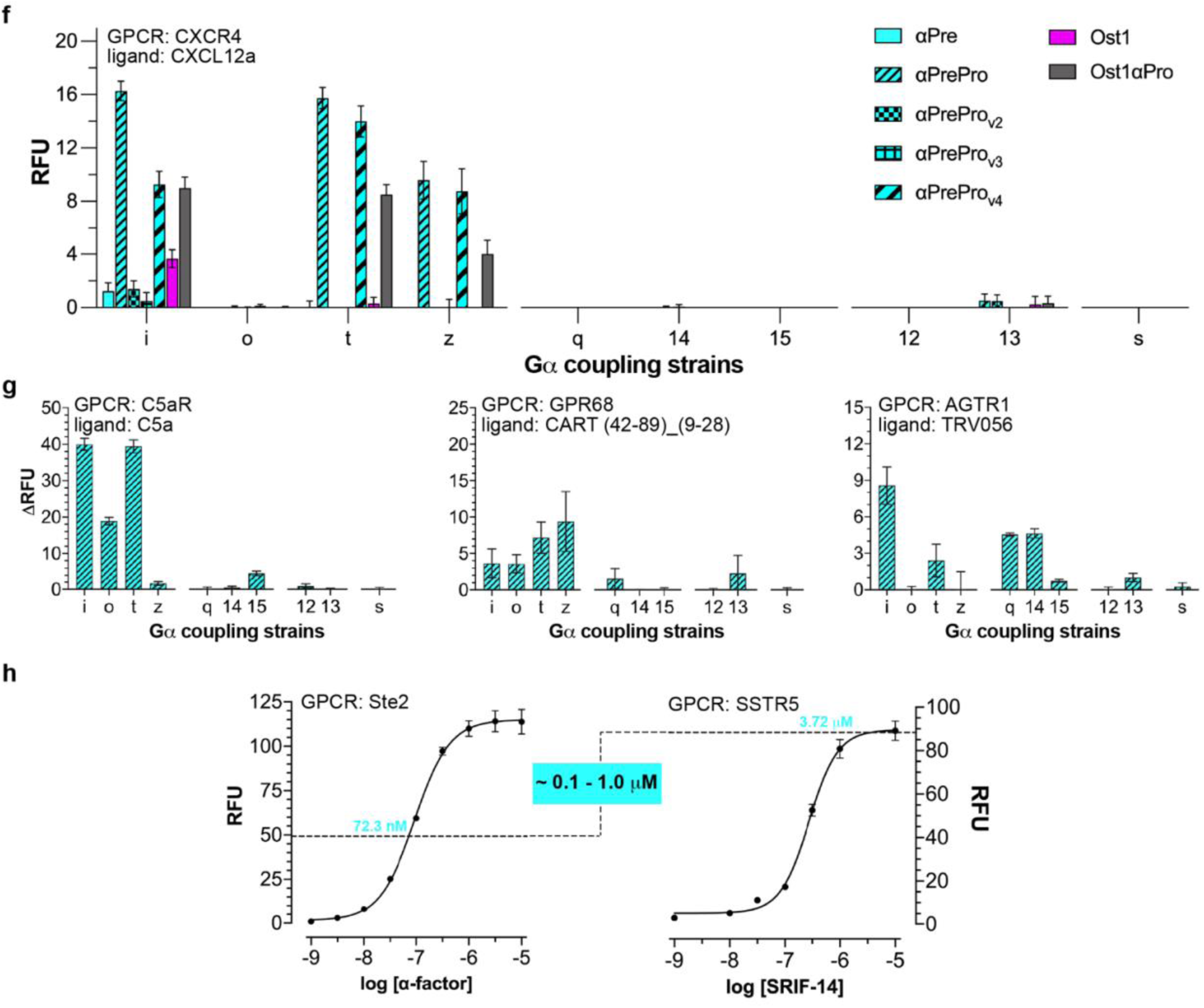
Characterization of secretion signals for PIONEER system design. **a**, Amino acid sequences of secretion signals selected for evaluation. **b**, Quantification of secreted mTq2 fluorescence (relative fluorescence units, RFU) to assess secretion signal output. “neg ctrl” indicates cells transformed with empty vector. Data are shown as mean ± s.d. of *n* = 8 biological replicates. **c**, Secretion efficiency reported as the percentage of mTq2 detected in the medium relative to total mTq2 (intracellular plus secreted). Data are shown as mean ± s.e.m. of *n* = 8 biological replicates. **d-f**, Autocrine activation of SSTR5 (**d**), AGTR1 (**e**), and CXCR4 (**f**) in response to secretion of their cognate ligands using the indicated secretion signals. **g**, Additional autocrine signaling profiles driven by ligands secreted via the αPrePro signal. Fluorescence is reported as ΔRFU relative to strains expressing the GPCR and secretion signal only (no encoded ligand). **h**, Estimation of ligand secretion levels produced with αPrePro. Titration curves for *Ste2* (left) and SSTR5 (right) using exogenous ligands were used to estimate concentrations of secreted αPrePro-ligands (dashed line) based on fluorescence output from Fig. 1f. SSTR5 experiments were conducted in the DCyFIR G_αi_ strain; *Ste2* experiments were performed in the parental strain with native G_α_ (*Gpa1*). Data in **d-h** are shown as mean ± s.d. of *n* = 4 biological replicates. Results from **b-f** were used to score secretion signal performance in mTq2 (**b-c**) and ligand-secretion (**d-f**) assays, as presented in Fig. 1f.

**Fig. S5.**
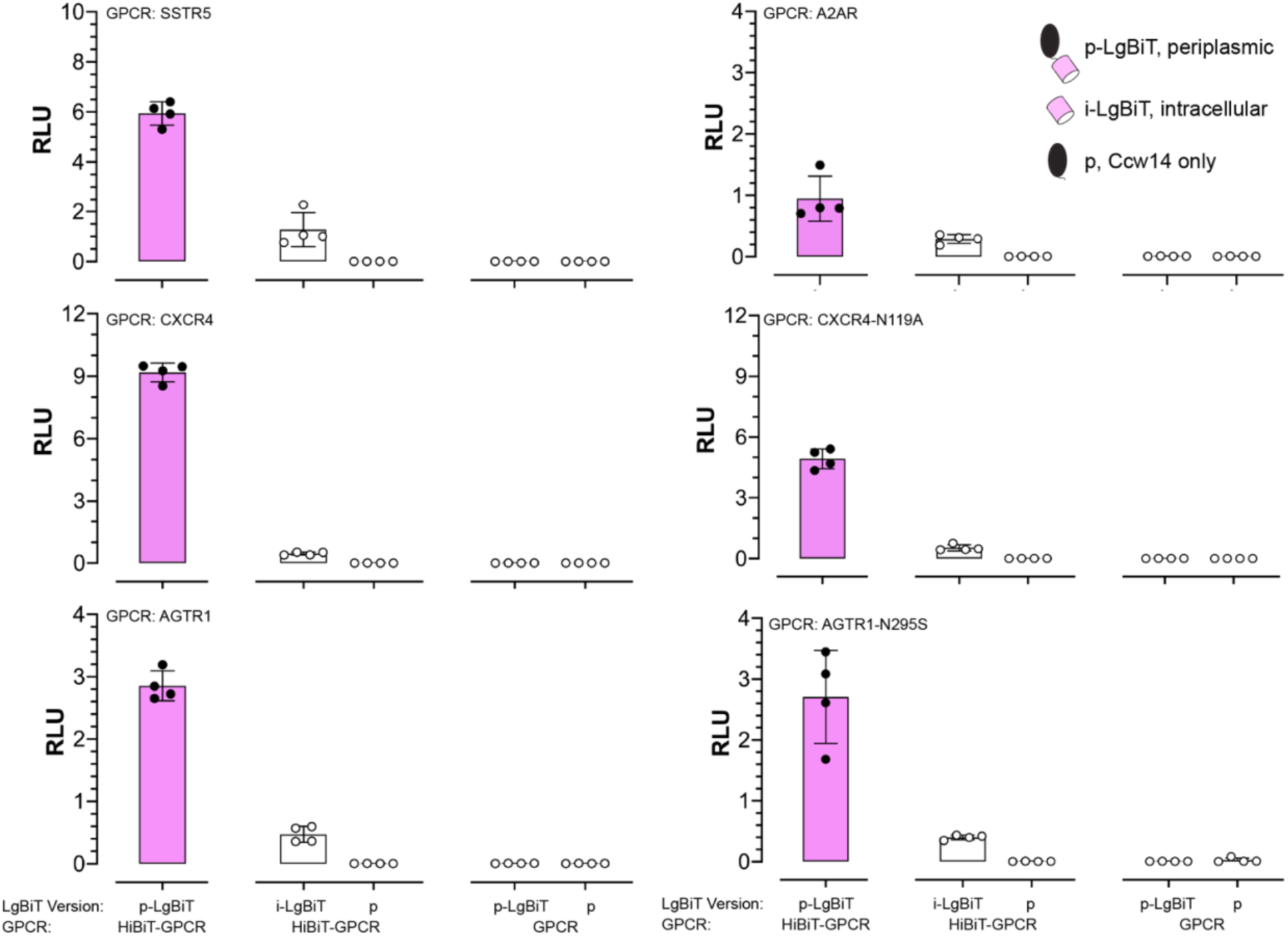
Control experiments for PIONEER-based detection of surface GPCRs. Control data corresponding to Fig. 2c. Luminescence is generated by reconstitution of NanoLuc between periplasmically displayed LgBiT and HiBiT-tagged GPCRs expressed at the cell surface. Luminescence is reported in relative luminescence units (RLU). Data represent mean ± s.d. of *n* = 4 biological replicates.

**Fig. S6.**
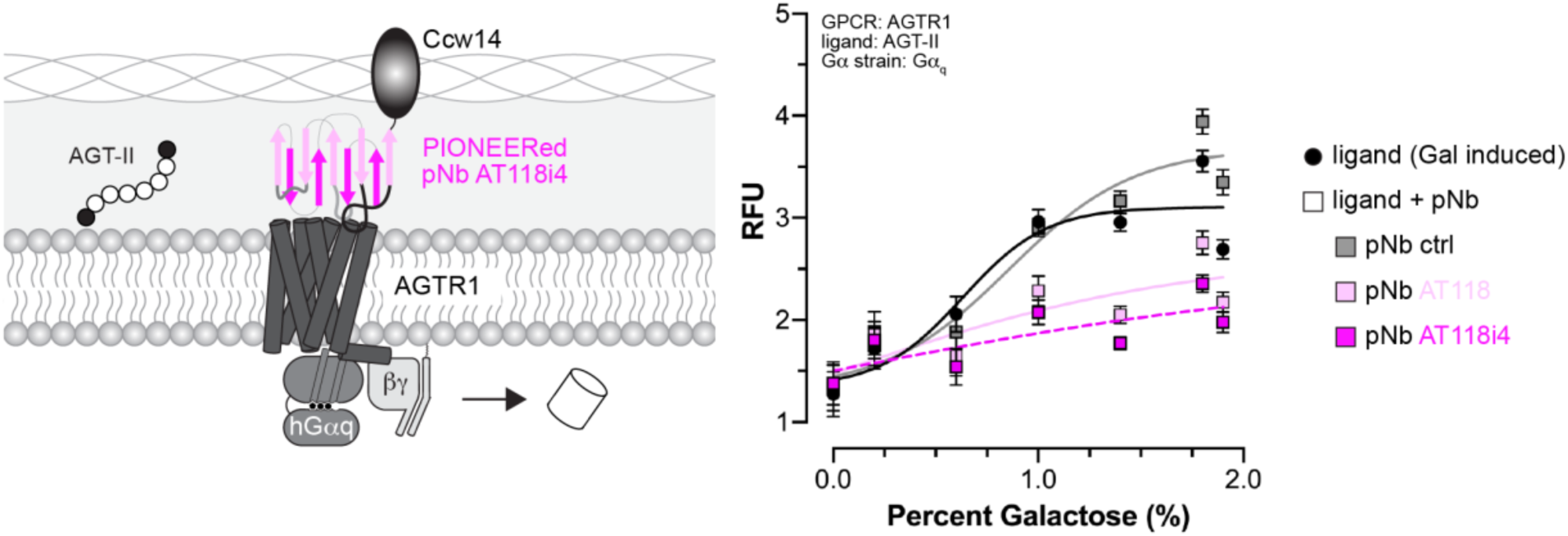
AGTR1-G_αq_ antagonism by PIONEERed nanobodies AT118 and AT118i4. Schematic and functional data showing inhibition of AGTR1-G_αq_ activation by angiotensin II (AGT-II) using PIONEERed nanobodies AT118 and AT118i4. This experiment parallels the G_αi_-based assays shown in Fig. 4b. Data represent mean ± s.d. of *n* = 4 biological replicates.

**Fig. S7.**
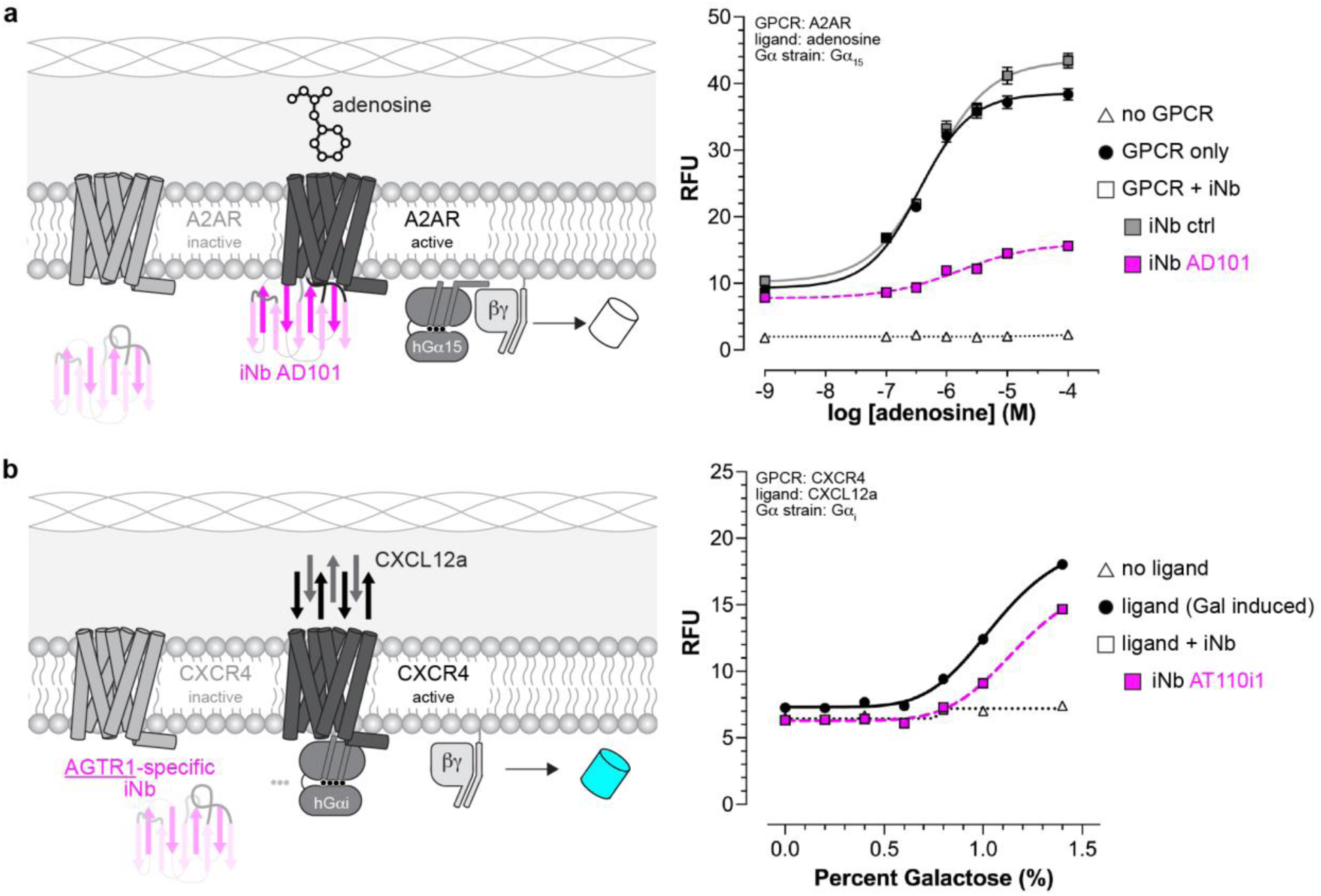
GPCR modulation by intracellular nanobodies. **a**, Schematic and functional data showing that intracellular nanobody AD101 reduces A2AR-G_α15_ signaling. This mirrors its inhibitory effect on A2AR-G_αi_ signaling shown in Fig. 5e. **b**, Schematic and data demonstrating that nanobody AT110i1, specific for AGTR1, does not enhance CXCR4-G_αi_ signaling in response to CXCL12a. This contrasts with the strong potentiation of AGTR1-G_αi_ signaling by AT110i1 shown in Fig. 5b. Data represent mean ± s.d. of *n* = 4 biological replicates.

